# Water and Electrolyte Homeostasis in a Mouse Model with Reduced ENaC Gamma Subunit Expression

**DOI:** 10.1101/2021.01.26.428168

**Authors:** Evan C. Ray, Alexa Jordahl, Allison Marciszyn, Aaliyah Winfrey, Tracey Lam, Yaacov Barak, Shaohu Sheng, Thomas R. Kleyman

**Affiliations:** Departments of Medicine, Pittsburgh, PA; Departments of Obstetrics, Gynecology & Reproductive Sciences, Pittsburgh, PA; Departments of Cell Biology, Pittsburgh, PA; Departments of Pharmacology and Chemical Biology, University of Pittsburgh, Pittsburgh, PA; Magee-Womens Research Institute, Pittsburgh, PA

**Keywords:** ENaC, Potassium Excretion, Salt Sensitivity, Extracellular Fluid Volume, Quantitative magnetic resonance

## Abstract

The epithelial Na^+^ channel (ENaC) promotes the absorption of Na^+^ in the aldosterone-sensitive distal nephron, colon, and respiratory epithelia. Deletion of genes encoding ENaC’s subunits results in early post-natal mortality. We present initial characterization of a mouse with dramatically suppressed expression of the γ subunit. We use this hypomorphic (γ^mt^) allele to explore the importance of ENaC’s γ subunit in homeostasis of electrolytes and body fluid volume. At baseline, γ subunit expression in γ^mt/mt^ mice is markedly suppressed in kidney and lung, while electrolytes resemble those of littermate controls. Challenge with a high K^+^ diet does not cause significant differences in blood K^+^, but provokes higher aldosterone in γ^mt/mt^ mice than controls. Quantitative magnetic resonance (QMR) measurement of body composition reveals similar baseline body water, lean tissue mass, and fat tissue mass in γ^mt/mt^ mice and controls. Surprisingly, euvolemia is sustained without significant changes in aldosterone or atrial natriuretic peptide. γ^mt/mt^ mice exhibit a more rapid decline in body water and lean tissue mass in response to a low Na^+^ diet than controls. Replacement of drinking water with 2% saline induces dramatic increases in body fat in both genotypes, and a selective transient increase in body water and lean tissue mass in γ^mt/mt^ mice. While ENaC in renal tubules and colon work to prevent extracellular fluid volume depletion, our observations suggest that ENaC in non-epithelial tissues may have a role in preventing extracellular fluid volume overload.

## Introduction

The epithelial Na^+^ channel (ENaC) has a key role in regulation of extracellular fluid volume and blood pressure, and renal K^+^ secretion. ENaC loss of function mutations in humans cause pseudohypoaldosteronism type 1 (PHA1), associated with urinary Na^+^ wasting, polyuria, hypotension, hyperkalemia and elevated aldosterone, reflecting extracellular fluid volume contraction (37). Gain-of-function mutations cause Liddle syndrome, characterized by hypertension, suppressed aldosterone, and hypokalemia secondary to enhanced renal tubular Na^+^ absorption and K^+^ secretion (37). The importance of ENaC in regulating Na^+^ handling and extracellular fluid volume suggest that ENaC activity may play a role in salt-sensitive hypertension (16, 25, 28).

The essential nature of the genes encoding ENaC’s subunits, α, β, or γ, has impaired investigation of the functional contributions of these subunits. A hypomorphic β subunit allele has proven useful in assessing the physiologic importance of this subunit, though complicated by inclusion of a Liddle syndrome mutation in the hypomorphic β subunit allele (29). Mice expressing this allele demonstrate more than twenty-fold reduction in β subunit expression but survive to adulthood, exhibiting a PHA1-like phenotype. Subsequent studies capitalized on this useful genetic tool to demonstrate the importance of the β subunit in modulation of respiratory fluid clearance, arterial myogenic vasoconstriction, renal vascular blood flow, and baroreception (9, 11, 14, 31).

Here, we present initial characterization of a gene-targeted mouse strain expressing a hypomorphic γ subunit allele, resulting in dramatically reduced expression in both kidney and lung but supporting viability in the homozygous state (γ^mt/mt^). This hypomorphic allele allows exploration of the importance of ENaC’s γ subunit in homeostasis of electrolytes and body fluid volume.

## Methods

### Generation and care of γ subunit hypomorphic mice

Gene-targeted mice were produced using homologous recombination in embryonic stem (ES) cells. *Scnn1g* exon 2 and 7 to 8 kB of flanking DNA were cloned from a Chromosome 7 bacterial artificial chromosome (BACPAC Genomics). The cDNA for the only known transcript of *Scnn1g* was inserted, in frame, distal to the start codon in *Scnn1g* exon 2 (Figure 1). An accompanying neomycin cassette decreased expression of the targeted locus. (27) Embryonic stem cells were electroporated with this construct. Correct insertion of the transgene was confirmed in G418-resistant stem cells by Southern blot. Chimeric mice carrying the targeted allele were produced at Charles River Inc. by microinjection of blastocysts with correctly targeted ES cell clones. Male progeny with germline incorporation of the transgene were selected as founders of independent mouse lines. Progeny were back-crossed for four to five generations into the 129S2/SvPasCrl (129sv, Charles River) or C57B/6 (Jackson Labs) background. Heterozygous (γ^+/mt^) mice from the two separate lines were cross-bred to generate experimental mice and littermate controls. Genotyping was performed in a single reaction with the following three primers: pER31 (intron 1 common forward primer: ACC TTA CTT GGC TCC TCT GTC CCT TC), pER32 (transgenic exon 2-3 junction reverse primer: GGA GGT CAC TCA CAG CAC TGT ACT TGT AG), and pER35 (wild-type intron 2 reverse primer: GGA GGC AGA TGC TAA CCT CAT TTC AGG). Amplification products were 633 bp (γ^+^) and 400 bp (γ^mt^). All control mice were littermates of γ^mt/mt^ mice.

**Figure 1.**
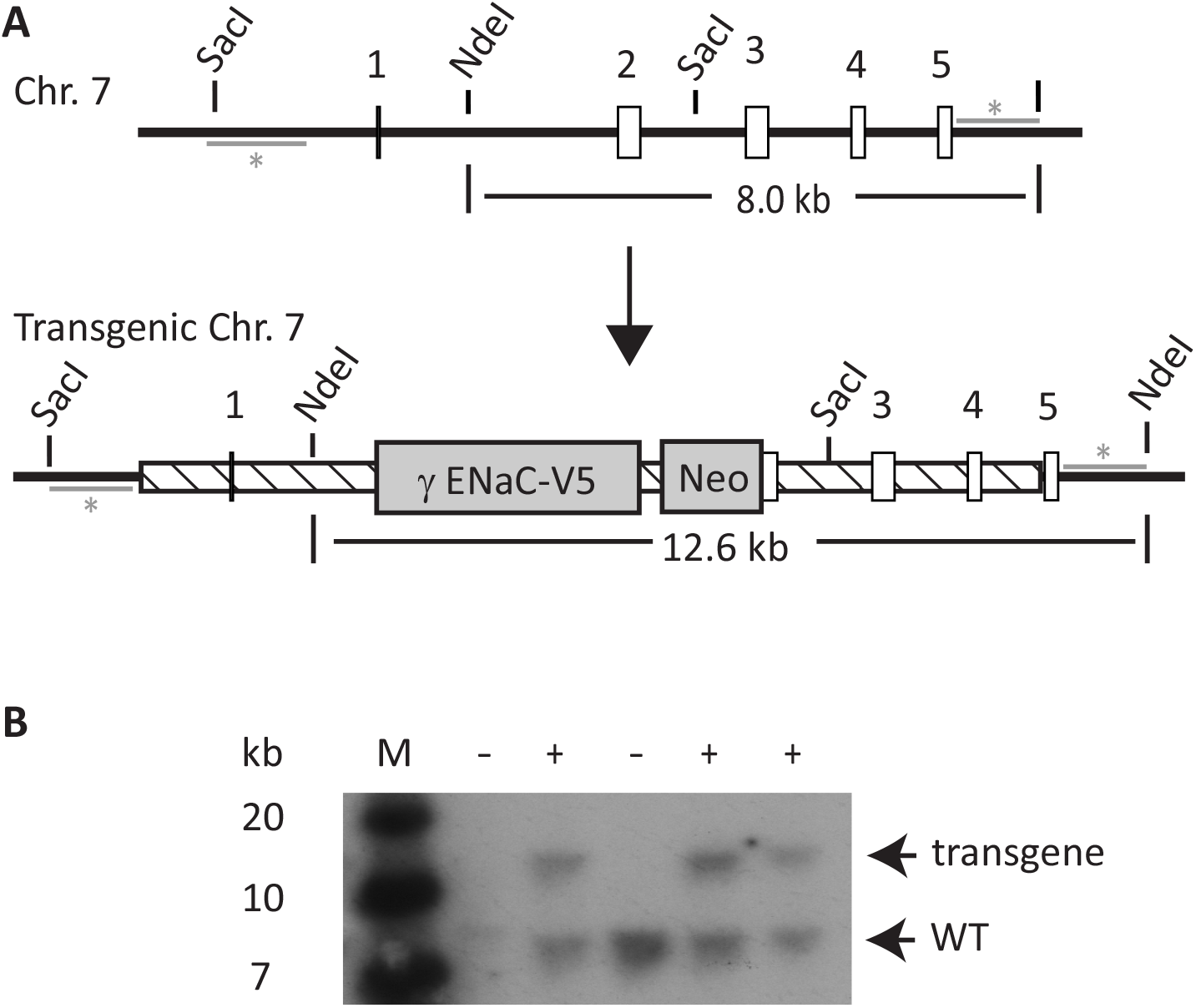
Generation of an *Scnn1g* Hypomorphic Allele. A schematic of the wild-type and transgenic *Scnn1g* locus (encoding the ENaC γ subunit) is shown in **A**. The genomic region corresponding to the targeting construct is represented as a hatched box. Open, boxes represent exons. Gray bars (with *) represent radiolabeled probes used for Southern blot. These are external to the targeting construct. Neo: neomycin-resistance cassette. γ ENaC-V5: cDNA for ENaC’s γ subunit, with C-terminal V5 tag, driven in frame by the start codon of the native gene. Restriction sites used for Southern blot are marked. **B**. Homologous recombination of the 5’ arm was analyzed by Southern blot of SacI-digested ES cell genomic DNA with the 5’ probe (not shown). Shown is a Southern blot with the 3’ probe of NdeI-digested genomic DNA of three representative clones from the 5’ analysis, confirming homologous recombination of the 3’ arm, which manifests as a shift in the size of the probed fragment from 8.0 to 12.6 kb. *+*: clones with correct homologous recombination of the 3’ arm. M: KB markers.

Regular diet included 0.94% K^+^ and 0.23% Na^+^ (Prolab Isopro RMH 3000, LabDiet). High K^+^ diet contained 5.2% K^+^ (as KCl) and 0.3% Na^+^ (Teklad TD.09075, Envigo). Low Na^+^ diet (TD.90228) contained 0.01-0.02% Na^+^ and 0.8% K^+^. 2% saline was unbuffered 2 g/dL NaCl in dH_2_0 (w/v). All animal work was approved by University of Pittsburgh Institutional Animal Care and Use Committee.

### Measurement of electrolytes and hormones

Whole blood was collected upon mouse sacrifice via aspiration from a cardiac ventricle and measurements were performed immediately using an iSTAT handheld device (Abbott Point of Care, Inc.). ELISAs for aldosterone (Enzo Life Sciences, catalog number ADI-900-173) or ANP (Sigma-Aldrich catalog number RAB00385) were performed on plasma diluted 1:25 to 1:200 or 1:4, respectively.

### Immunoblotting

Tissues were stored at -80 degrees C° until use. Approximately 50 mg of tissue was homogenized using a Dounce homogenizer in 0.15 mL (for kidney) or 0.2 mL (for lung) HEPES-buffered saline plus Phosphatase Inhibitor Cocktail Set II and Protease Inhibitor Cocktail Set III (Calbiochem). Protein was measured using a Pierce™ BCA protein assay kit (Thermo Fisher) and mixed 1:1 with 2x Laemmli buffer (Bio-Rad) plus 5% β-mercaptoethanol. Sixty μg of protein were loaded per well in a 4-15% acrylamide TGX gel (Bio-Rad) and run in 1x Tris-Glycine SDS buffer. Proteins were transferred to nitrocellulose, and blocked in 10% milk in TBST. Primary antibodies included anti-γ subunit (StressMarq; catalogue number SPC-405; 0.5 μg/mL), anti-β subunit (StressMarq; catalogue number SMC-241; 1 μg/mL), and anti-GAPDH (ProteinTech, 0.1 μg/mL). After incubation with appropriate HRP-linked secondary antibodies, blots were digitally analyzed a ChemiDoc Imaging System (Bio-Rad).

### Body composition analysis

*In vivo* mouse body composition was measured by quantitative magnetic resonance using a 100H Body Composition Analyzer (EchoMRI™)(40, 43). QMR was recently shown to be effective at detecting progressive differences in body water in response to changes in dietary Na^+^ and water or mineralocorticoid treatment (24). Measurements were performed in late afternoon. Tag-free, un-anesthetized, ∼10 week-old mice were weighed, immediately placed in a restraint tube (EchoMRI), and inserted into the analyzer. The tube was carefully examined to ensure absence of urine or other fluid from previously measured mice. Choice of restraint tube influenced measurements, therefore the same tube was used for all measurements. Each measurement required less than 3 minutes. After measurement, each mouse was returned immediately to its cage. After all mice were measured once, the all mice were individually weighed and measured again. Data shown represent mean of two measurements. Mouse weights and body water declined by 0.2 to 0.6 g between measurements (∼1 hour apart), so mice were measured in the same order each session.

### Statistics

Outliers were removed using the ROUT method (ROUT coefficient of 1%). Pair-wise comparisons were performed by Student’s *t-*test (α = 0.05). For multiple, independent comparisons, *p* was adjusted using Sidak’s method (38). For non-independent, repeated measurements (body composition over time), false discovery rates were controlled using Benjamini, Krieger, and Yekutieli’s two-stage step-up method (Q = 0.05), and *p* values adjusted accordingly (3). Analyses were performed using Prism 8.4.2 (GraphPad). Reported errors represent standard deviation.

## RESULTS

Immunoblots of whole kidneys and lungs from transgenic mice (γ^mt/mt^) and littermate controls (γ^+/+^) revealed reduced γ subunit expression (Figure 2). In kidney, γ subunit (normalized to γ^+/+^) was reduced from 100 ± 10 % (N = 5) to 8 ± 4 % (N = 6; p < 0.0001). In lung, it was reduced from 100 ± 43 % (N = 6) to 16 ± 4% (N = 4; p < 0.01). β subunit expression was not significantly different: 100 ± 60 % (N = 5) in control kidneys versus 62 ± 21 % (N = 6, p = NS) in γ^mt/mt^ kidneys, and 100 ± 49 % (N = 6) in control lungs versus 144% ± 132 % (N = 5) in γ^mt/mt^ lungs (p = NS).

**Figure 2.**
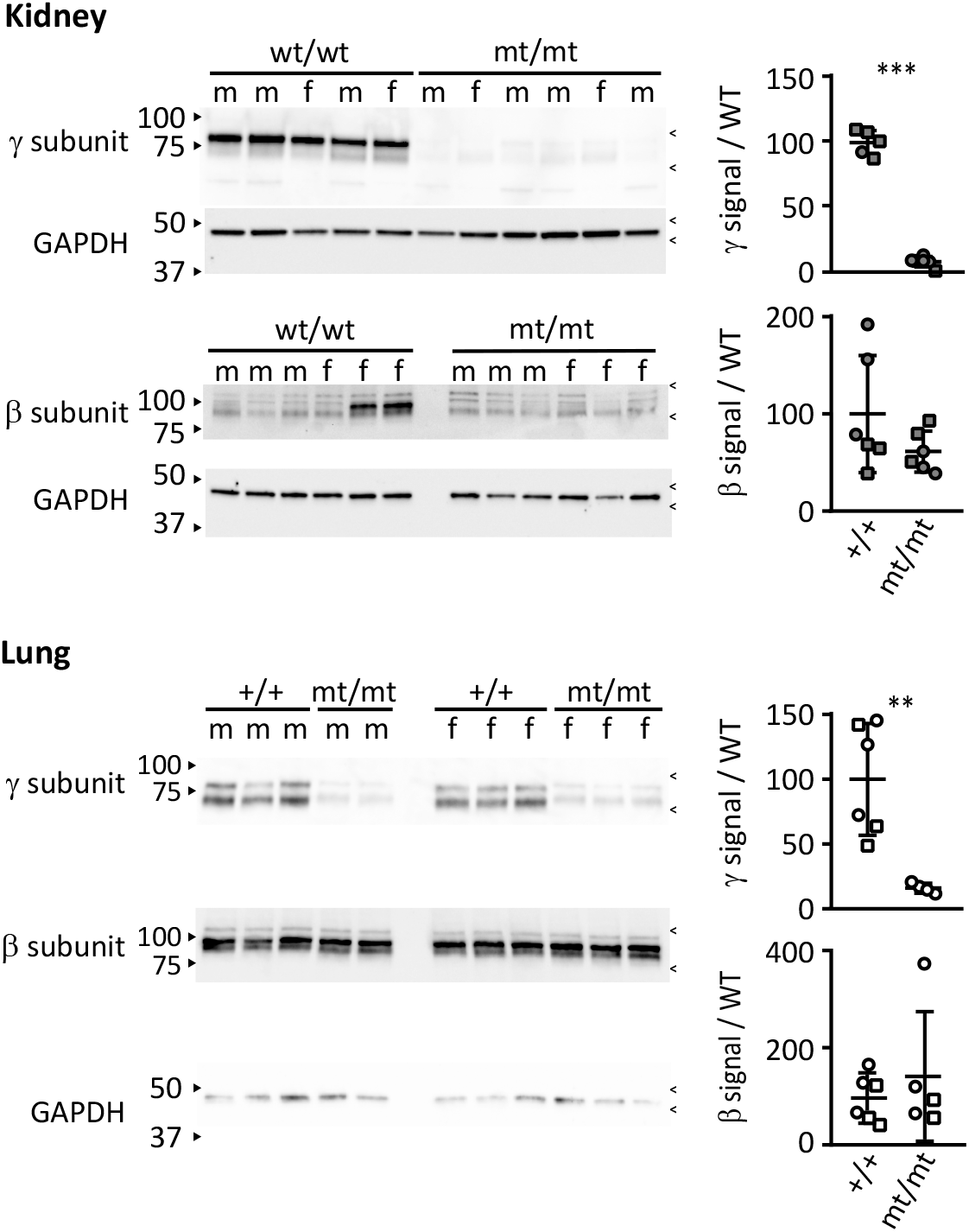
γ^mt/mt^mice express significantly less γ subunit in kidney and lung. Immunoblots of kidney and lung tissue lysates were probed with antibodies against the β or γ subunit of ENaC, then stripped and probed with antibodies against GAPDH. Arrowheads on the left of each blot show molecular weight marker positions. Carrots on the right of each blot indicate approximate region taken for signal quantification. Dot plots show protein normalized to mean signal from γ^+/+^ mice. Squares represent males, circles represent females. Open or shaded symbols represent data from 129sv or C57BL/6 mice, respectively. γ subunit protein levels in γ^mt/mt^ mouse kidneys were reduced from 100 ± 10 % (mean ± SD, N = 5 mice) to 8 ± 4 % (N = 6). ***: p < 0.0001 (two-tailed Student’s t-test). In lung, the γ subunit was reduced from 100 ± 43 % (N = 6) to 16 ± 4% (N = 4) **: p < 0.01 (two-tailed Student’s t-test). The kidney β subunit point estimate decreased from 100 ± 60 % (N = 6) to 62 ± 9% (N = 6), but the difference was not significant. Lung β subunit protein signal also did not differ significantly between groups: 100 ± 49 % (N = 6) for γ^+/+^ vs. 144 ± 132 % for γ^mt/mt^ (N = 5).

**Figure 3.**
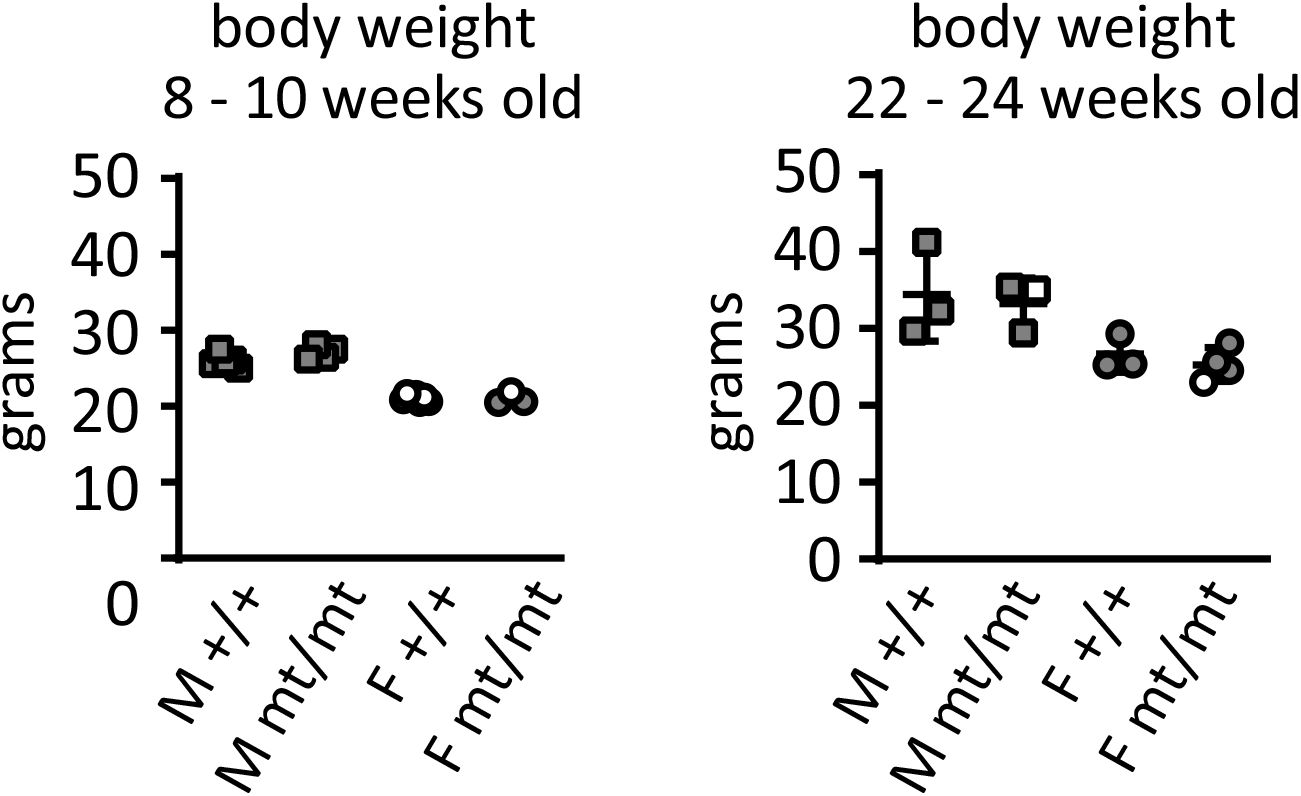
Pups born to γ^+/mt^parents exhibit genotype frequencies consistent with Mendelian inheritance and gain weight normally. Left and right plots demonstrate that γ^mt/mt^ mice exhibit weights similar to age- and sex-matched γ^+/+^ littermate controls. 8 to 10 week-old mice: γ^+/+^ males weighed 26.0 ± 0.9 g, N = 5; γ^mt/mt^ males weighed 27.1 ± 0.9 g, N = 4, p = NS. γ^+/+^ females weighed 21.1 ± 0.4 g, N = 7; γ^mt/mt^ females weighed 21.0 ± 0.8 g, N = 3, p = NS. 22 to 24 week-old mice: γ^+/+^ males weighed 34.4 ± 6.1 g, N = 3; γ^mt/mt^ males weighed 33.3 ± 3.3 g, N = 3, p = NS; γ^+/+^ females weighed 26.7 ± 2.3 g, N = 3; γ^mt/mt^ females weighed 25.35 ± 2.2 g, N = 4, p = NS. (Comparisons examined using two-tailed Student’s t-test.) Open or shaded symbols represent data from 129sv or C57BL/6 mice, respectively.

Because deletions of ENaC subunits exhibit early post-natal mortality, we examined litter size and genotype ratios in γ^+/mt^ crosses. (2) Litter size at weaning was 4.4 ± 1.8 for 129sv pups (25 litters) and 5.2 ± 1.9 for C57BL/6 pups (5 litters). These resemble previously reported wild-type litter sizes of 4.6 for 129sv mice and 5.5 for C57BL/6 mice (12). Analysis of litters from 23 heterozygote crosses in the 129sv background revealed that live γ^mt/mt^ pups were present at near-Mendelian ratios (29 out of 103 compared to 28/103 and 46/103 for γ^+/+^ and γ^+/+^ pups, respectively). Weights of γ^mt/mt^ mice resembled age- and sex-matched controls. These data indicate no significant pre- or peri-natal mortality or abnormal growth of γ^mt/mt^ mice.

Pharmacologic or genetic inhibition of ENaC causes hyperkalemia (2, 32, 44). On a regular diet, blood K^+^, Na^+^, Cl^-^, total CO_2_ (tCO_2_), urea nitrogen (BUN), and hemoglobin (Hb) from γ^mt/mt^ mice resembled controls (Table 1 and Figure 4). Measurements in 129sv mice resembled C57B6/J mice. Blood K^+^ also did not differ between genotypes on a high K^+^ diet (Table 2 and Figure 5). The point-estimate for blood K^+^ in γ^mt/mt^ males exceeded that of γ^+/+^ males, but did not reach significance (adjusted p = 0.09). Blood Na^+^ was lower in γ^mt/mt^ females than in γ^+/+^ females (adjusted p = 0.04).

**Table 1.** Blood measurements on a regular diet.

| | $\gamma^{+/+}$ | $\gamma^{mt/mt}$ | Male<br>$\gamma^{+/+}$ | Male<br>$\gamma^{mt/mt}$ | Female<br>$\gamma^{+/+}$ | Female<br>$\gamma^{mt/mt}$ |
| --- | --- | --- | --- | --- | --- | --- |
| Na<sup>+</sup> (mmol/L) | 145 ± 2 (21) | 144 ± 2 (16) | 145 ± 1 (13) | 145 ± 2 (10) | 144 ± 2 (8) | 143 ± 1 (6) |
| K<sup>+</sup> (mmol/L) | 4.9 ± 0.7 (20) | 4.9 ± 0.6 (16) | 5.0 ± 0.7 (12) | 4.9 ± 0.4 (10) | 4.7 ± 0.7 (8) | 4.8 ± 0.8 (6) |
| Cl<sup>-</sup> (mmol/L) | 113 ± 2 (14) | 113 ± 3 (16) | 113 ± 2 (9) | 113 ± 2 (10) | 114 ± 2 (5) | 112 ± 3 (6) |
| tCO<sub>2</sub> (mmol/L) | 24.8 ± 3.3 (20) | 24.5 ± 2.3 (16) | 26.5 ± 3.1 (12) | 25.2 ± 1.6 (10) | 22.3 ± 1.7 (8) | 23.3 ± 2.9 (6) |
| Urea Nitrogen (mg/dL) | 27 ± 5 (14) | 28 ± 5 (16) | 28 ± 5 (9) | 28 ± 5 (10) | 25 ± 5.5 (5) | 27 ± 6 (6) |
| Hemoglobin (mg/dL) | 13.1 ± 0.9 (20) | 13.6 ± 0.8 (16) | 13.3 ± 0.5 (12) | 13.5 ± 0.8 (10) | 12.9 ± 1.3 (8) | 13.8 ± 1.0 (6) |
| Creatinine (mg/dL) | < 0.2 md/dL | < 0.2 md/dL | < 0.2 md/dL | < 0.2 md/dL | < 0.2 md/dL | < 0.2 md/dL |
| Aldosterone (pg/mL) | 590 ± 720 (11) | 880 ± 610 (9) | 420 ± 290 (8) | 880 ± 580 (5) | 1040 ± 1380 (3) | 890 ± 750 (4) |
| ANP (pg/mL) | 221 ± 114 (11) | 191 ± 101 (10) | 259 ± 112 (8) | 227 ± 104 (6) | 121 ± 11 (3) | 136 ± 77 (4) |

Blood parameters from mice on a regular diet are shown: mean ± standard deviation (N). Electrolytes, urea nitrogen, hemoglobin, and creatinine were assayed in whole blood. All creatinine values were below the detection threshold. Aldosterone and atrial natriuretic peptide (ANP) measurements were measured in plasma. There were no significant differences between $\gamma^{+/+}$ mice and $\gamma^{mt/mt}$ mice, either in male mice or female mice, or when mice from both sexes were pooled, as determined by two-tailed Student's t-test with adjustment of *p* values for multiple comparisons (two sexes) using Sidak's method.

**Table 2.** Blood measurements on a high K^+^ diet.

| | $\gamma^{+/+}$ | $\gamma^{mt/mt}$ | Male<br>$\gamma^{+/+}$ | Male<br>$\gamma^{mt/mt}$ | Female<br>$\gamma^{+/+}$ | Female<br>$\gamma^{mt/mt}$ |
| --- | --- | --- | --- | --- | --- | --- |
| Na<sup>+</sup> (mmol/L) | 147 ± 2 (19) | 147 ± 3 (24) | 147 ± 2 (10) | 148 ± 2 (17) | 146 ± 2 (9) | 144 ± 2 (7)* |
| K<sup>+</sup> (mmol/L) | 5.2 ± 0.6 (19) | 5.4 ± 0.7 (24) | 5.0 ± 0.7 (10) | 5.5 ± 0.7 (17) | 5.5 ± 0.5 (9) | 5.2 ± 0.7 (7) |
| Cl<sup>-</sup> (mmol/L) | 119 ± 3 (19) | 119 ± 3 (23) | 118 ± 3 (10) | 119 ± 3 (16) | 119 ± 4 (9) | 119 ± 4 (7) |
| tCO<sub>2</sub> (mmol/L) | 20 ± 2 (19) | 20 ± 2 (24) | 21 ± 2 (10) | 21 ± 1 (17) | 20 ± 2 (9) | 18 ± 2 (7) |
| Urea nitrogen (mg/dL) | 26 ± 4 (19) | 28 ± 4 (24) | 28 ± 4 (10) | 28 ± 4 (17) | 25 ± 3 (9) | 29 ± 6 (7) |
| Hemoglobin (mg/dL) | 13.3 ± 0.9 (19) | 13.5 ± 0.9 (24) | 13.4 ± 0.7 (10) | 13.6 ± 0.8 (17) | 13.2 ± 1.1 (9) | 13.4 ± 1 (7) |
| Creatinine (mg/dL) | < 0.2 md/dL | < 0.2 md/dL | < 0.2 md/dL | < 0.2 md/dL | < 0.2 md/dL | < 0.2 md/dL |
| Aldosterone (pg/mL) | 3100 ± 2500 (19) | 4900 ± 2500 (23)* | 3100 ± 2800 (10) | 4600 ± 2500 (17) | 3000 ± 2200 (9) | 5800 ± 2700 (6) |

Animals were given high K<sup>+</sup> diet (5.2% K<sup>+</sup>, as KCl) for 10 days. Electrolytes, urea nitrogen, hemoglobin, and creatinine were assayed in whole blood. Creatinine values were all below the detection threshold. Aldosterone was measured in plasma. Differences were assessed using a two-tailed Student's t-test, with adjustment of *p* values for multiple comparisons (two sexes) using Sidak's method. \*: *p* < 0.05 for difference between $\gamma^{+/+}$ and $\gamma^{-/-}$ .

**Figure 4.**
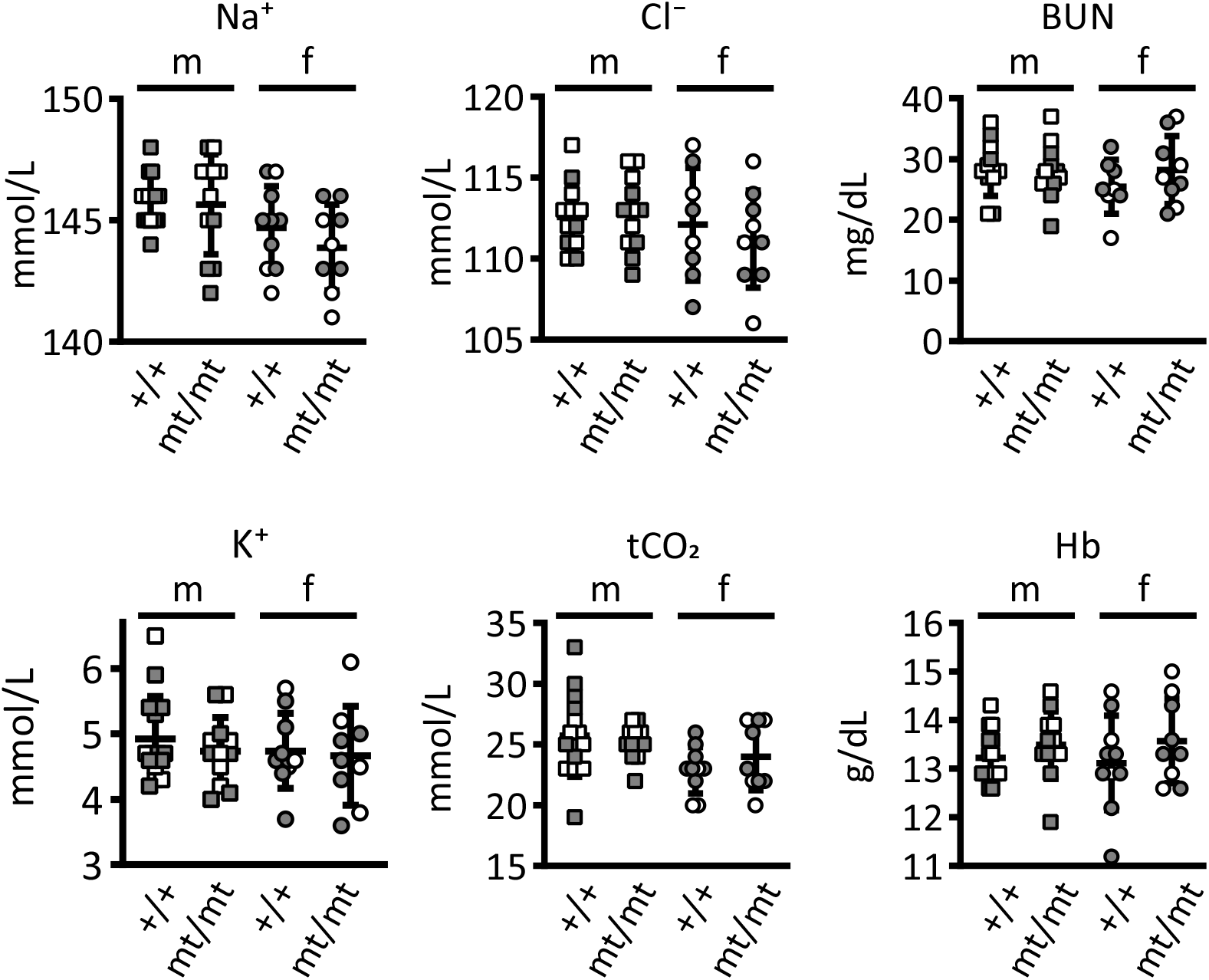
Blood analytes are similar in γ^mt/mt^and γ^+/+^mice. Blood parameters from mice on a regular diet are shown. Electrolytes, urea nitrogen, hemoglobin, and creatinine were measured in whole blood. Open symbols represent mice in the 129sv background; shaded symbols represent the C57BL/6 background. There were no significant differences between γ^+/+^ mice and γ^mt/mt^ mice, as determined by two-tailed Student’s t-test with adjustment of *p* values for multiple comparisons (two sexes) using Sidak’s method. Please see Table 1 for means and standard deviations.

**Figure 5.**
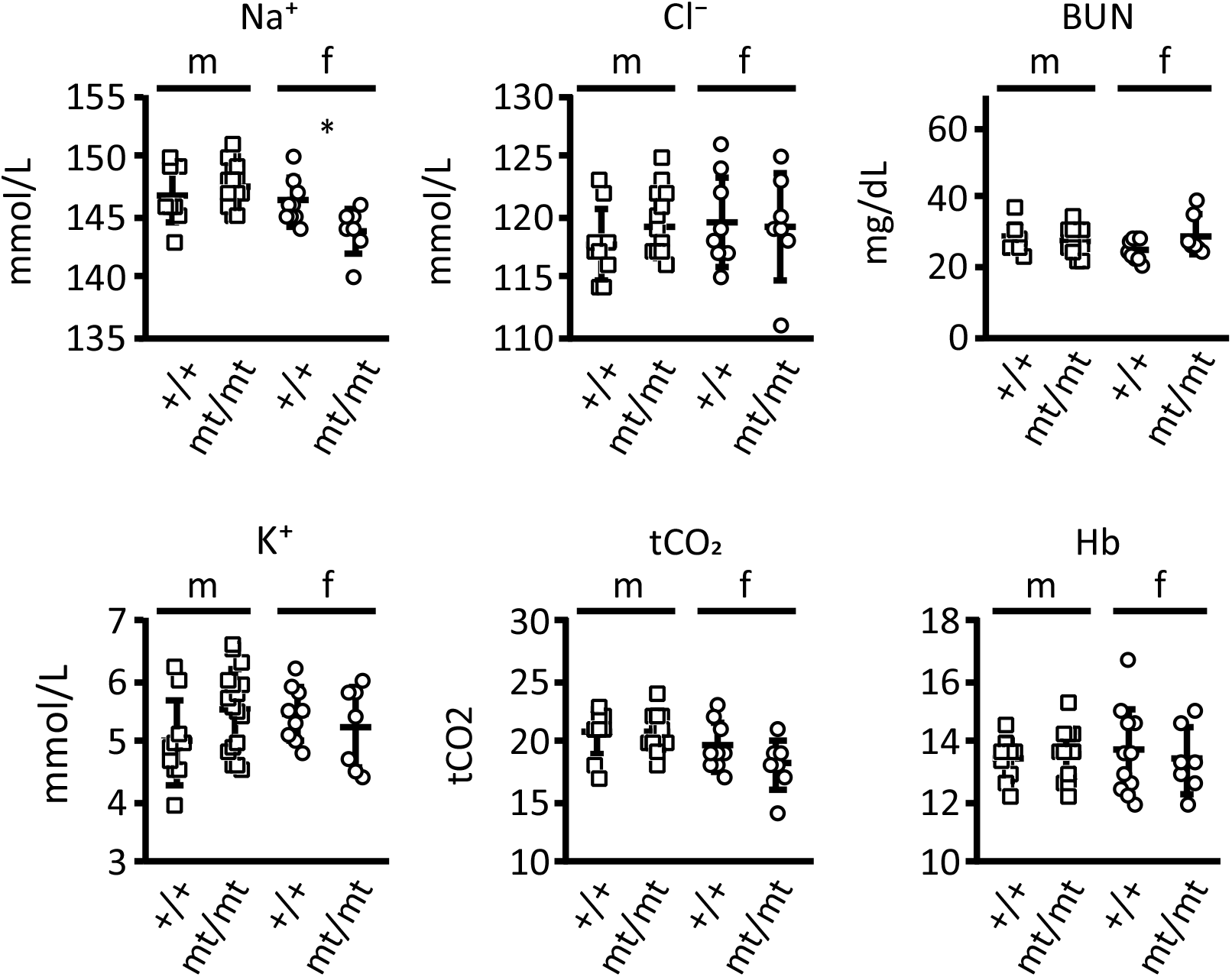
On a high K^+^diet, blood electrolyte, BUN and Hb concentrations in γ^mt/mt^ mice largely resemble those of γ^+/+^mice. 129sv mice were provided with 5.2% K^+^ (as KCl) diet for 10 days before blood collection. Electrolytes, urea nitrogen, hemoglobin, and creatinine were measured in whole blood. Blood Na^+^ was higher in female γ^+/+^ mice than in γ^mt/mt^ mice (146 ± 2, N = 9 vs. 144 ± 2, N = 7; adjusted p = < 0.05). There were no other significant differences between γ^+/+^ mice and γ^mt/mt^ mice. Groups were compared using two-tailed Student’s t-test with adjustment of *p* values for multiple comparisons in two sexes using Sidak’s method. See Table 2 for means and standard deviations.

Increased blood K^+^ stimulates aldosterone secretion, as does decreased blood volume. Either of these could increase aldosterone in the plasma of γ^mt/mt^ mice. Elevated aldosterone enhances urinary K^+^ excretion and would attenuate any increase in blood K^+^ in γ^mt/mt^ mice. On a regular diet, plasma aldosterone from γ^mt/mt^ mice were similar to those of γ^+/+^ mice (γ^+/+^: 580 ± 720 pg/mL, N = 11; γ^mt/mt^: 880 ± 610, N = 9, p = NS) Atrial natriuretic peptide (ANP) levels were also similar between groups (Table 1). Mean plasma aldosterone levels increased in both γ^mt/mt^ and γ^+/+^ mice when mice were placed on a high K^+^ diet for 10 days, but they were higher in γ^mt/mt^ mice than in controls (Figure 6 and Table 2; γ^+/+^ 3100 ± 2500 pg/mL, N = 19; γ^mt/mt^ 4900 ± 2500, N = 23, p < 0.05). This difference was not evident when males and females were analyzed separately.

**Figure 6.**
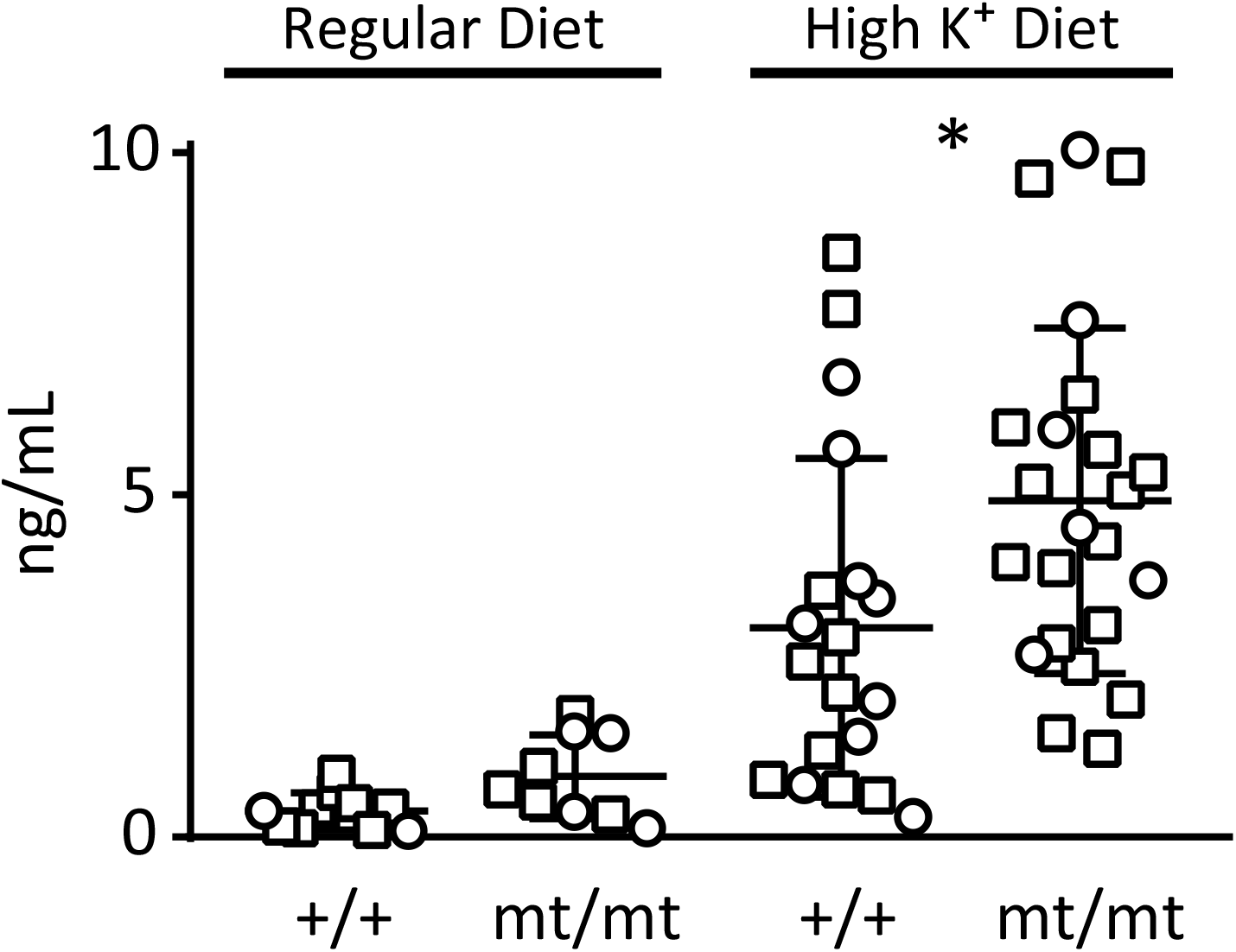
Plasma aldosterone levels in γ^mt/mt^mice are similar to controls at baseline, but are higher than γ^+/+^littermate controls on a high K^+^diet. 129sv mice were maintained on a regular (0.94% K^+^) diet, or transitioned to a 5.2% K^+^ (as KCl) diet for 10 days. Squares represent males, circles represent females. On a regular diet, there was no differences in plasma aldosterone between γ^+/+^ mice (590 ± 720 pg/mL, N = 11) and γ^mt/mt^ mice (880 ± 610 pg/mL, N = 9). On the high K^+^ diet, aldosterone increased in both groups (to 3100 ± 2500 pg/mL, N = 19 in γ^+/+^ mice, adjusted p < 0.05 compared to γ^+/+^ mice on a regular diet; and 4900 ± 2500 pg/mL, N = 23 in γ^mt/mt^ mice, adjusted p < 0.001 compared to regular diet). Aldosterone on the high K^+^ diet was higher in γ^mt/mt^ mice (* p =0.01, as determined by Student’s t-test, adjusted for multiple comparisons using Sidak’s multiple comparisons test.) Please see Table 2 for analysis by sex.

We hypothesized that γ^mt/mt^ mice would exhibit similar total body water and lean tissue mass when compared against γ^+/+^ littermates on a regular diet, given that plasma aldosterone and ANP levels were similar. Quantitative magnetic resonance revealed no significant differences in percent body water, lean tissue mass, or fat tissue mass (Figure 7). For γ^+/+^ mice, body water was 67.7 ± 2.7 % (N = 10; 1 female and 9 males), compared to 67.4 ± 2.6 % for γ^mt/mt^ mice (N = 7; 2 female and 5 males; p = NS). Lean tissue mass was 79.9 ± 2.5 % vs. 79.4 ± 3.0 % (p = NS), and fat composition was 12.0 ± 3.1 % vs. 12.8 ± 3.1 % (p = NS).

**Figure 7.**
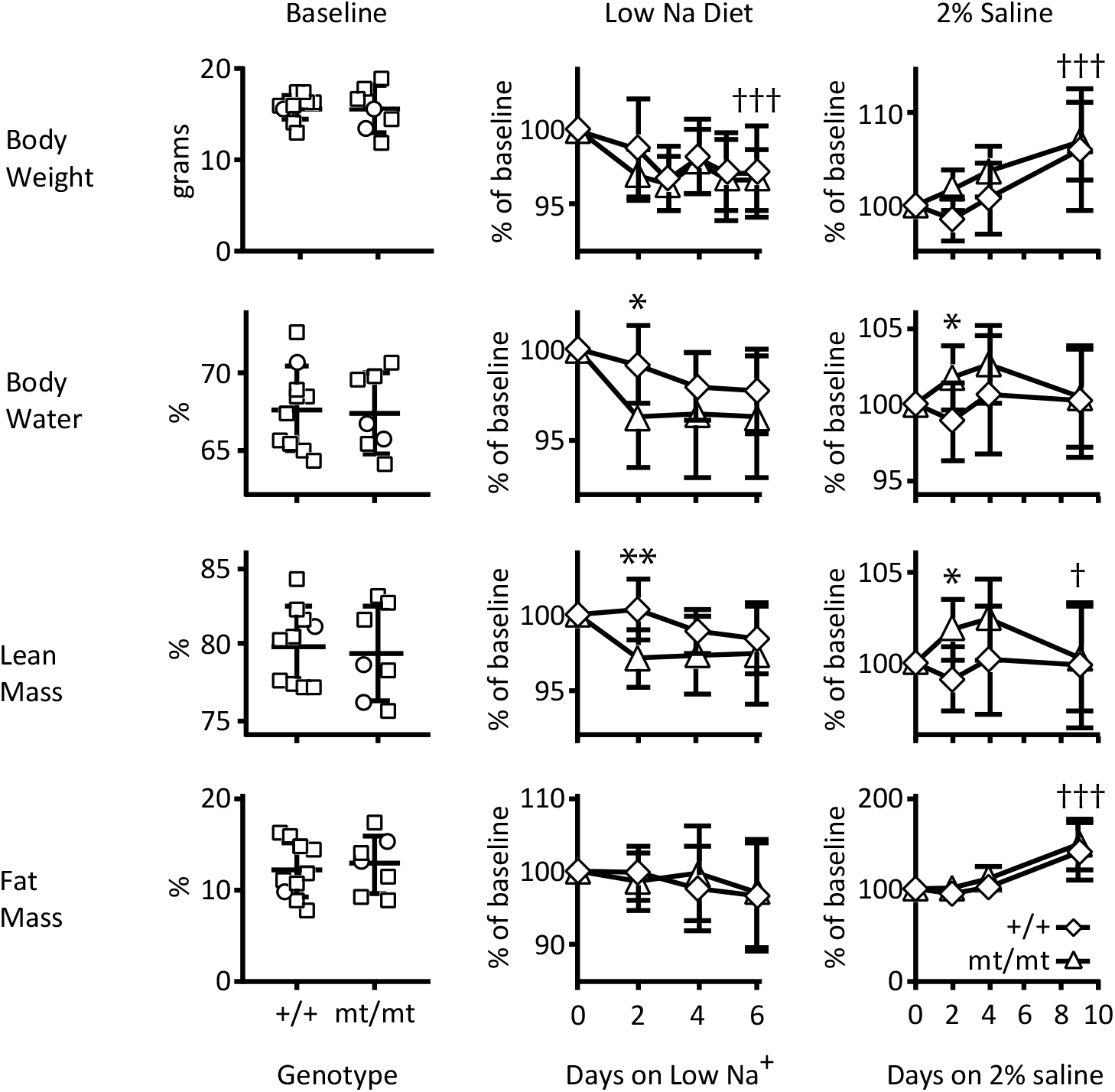
Body composition analysis reveals no significant difference in body water, lean tissue mass, or fat mass on a regular diet, but shows differences of body water and lean tissue mass to in response to a low or high salt diet. Body composition in 129sv mice was measured using quantitative magnetic resonance. Circles represent female mice, squares represent males. Starting weights were similar between groups. Prior to dietary manipulation, as a percentage of total body weight, there were no differences in body composition. These animals were placed on a Na^+^-depleted (0.01-0.02% Na^+^) diet, and subsequent body composition variables were normalized to baseline for each animal. Triangles and diamonds represent γ^mt/mt^ and γ^+/+^ mice, respectively. Weight declined but did not differ between genotypes. At day 2, normalized body water was lower in γ^mt/mt^ mice than in γ^+/+^ mice (p = 0.03), and normalized lean tissue mass was lower in γ^mt/mt^ mice (p < 0.01). Normalized fat content did not differ between γ^+/+^ and γ^mt/mt^ mice. Composition did not differ on subsequent days. After a 10 day recovery on regular diet, effect of 2% saline on body composition was evaluated. 2% saline increased weights overall, but genotypes did not differ. On day 2, normalized body water and lean tissue increased in γ^mt/mt^ mice compared to controls (p = 0.03 and p = 0.04, respectively). Thereafter, composition did not differ between genotypes. Body fat increased similarly in γ^+/+^ and γ^mt/mt^ mice. Baseline differences between genotypes were assessed by two-tailed t-test. Changes in body composition following dietary changes were analyzed using two-way ANOVA (†: < 0.05, †: < 0.01, †††: p < 0.0001 for time as a source of variation). Differences in genotypes over time were assessed by two-stage linear step-up procedure of Benjamini, Krieger and Yekutieli (*: p < 0.05. **: p < 0.01).

γ^mt/mt^ mice did not have evidence of volume depletion at baseline. To evaluate whether compensatory mechanisms could be overcome by physiologic stress, animals were given a Na^+^-depleted (0.01-0.02% Na^+^) diet (Figure 7). Over six days, body weights decreased significantly overall (p < 0.0001). However, genotype-dependent differences in body weight were not observed. Similarly, normalized body water percentage decreased over this period (P < 0.0001). The γ^mt/mt^ mice exhibited lower normalized body water at day 2 than controls (99.1 ± 2.1% for γ^+/+^ mice, N = 10; 96.3 ± 2.7 % for γ^mt/mt^ mice, N = 7; adjusted p = 0.03; Figure 7). Subsequently, differences between genotypes lost significance. Normalized lean tissue mass also decreased overall (p = 0.002). At day 2, γ^mt/mt^ mice exhibited lower normalized lean mass than controls (100.3 ± 2.1% for γ^+/+^ mice, N = 10; 97.1 ± 1.9 % for γ^mt/mt^ mice, N = 7; adjusted p < 0.01). Body fat did not change in either group.

The ability of ENaC to promote renal and gastrointestinal Na^+^ retention has led to the hypothesis that the channel may contribute to salt-sensitive hypertension (16, 34, 35). We therefore asked whether γ^mt/mt^ mice were protected from fluid volume overload during an obligatory increase in Na^+^ intake associated with replacement of electrolyte-free drinking water with 2% saline. Mice prefer saline to electrolyte-free water and consume more than 6 mL of 2% saline per day (19). The saline increased body weights significantly by 9 days, with no difference between groups (106 ± 6.8 %, N = 10 γ^+/+^ mice and 107 ± 4.2 %, N = 7 γ^mt/mt^ mice, on day 9, p = NS; Figure 7). Body water increased in γ^mt/mt^ mice relative to controls at day two (to 101.7 ± 2.1% vs. 98.9 ± 2.6%, respectively; adjusted p = 0.03). γ^mt/mt^ mice also exhibited higher lean tissue mass on day two (101.9 ± 1.7% vs. 99.1 ± 1.7%, respectively; p = 0.04). Body water and lean tissue masses did not differ significantly on later days. Body fat increased dramatically, but similarly, in γ^+/+^ and γ^mt/mt^ mice over 9 days (to 140.7 ± 30.7 % and 149.7 ± 28.3 %, respectively).

## Discussion

These results demonstrate a grossly healthy mouse model with reduced expression of ENaC’s γ subunit. Homozygous mice have no evidence of volume-depletion or impaired K^+^ handling at baseline, but exhibit aldosterone on a high K^+^ diet suggesting impaired K^+^ secretion. They experience enhanced sensitivity of body fluid volume to changes in dietary Na^+^, with more rapid loss of fluid and lean tissue on a low Na^+^ diet. Surprisingly, a transient larger increase in fluid and lean tissue mass was observed when water was replaced with 2% saline to drink. A dramatic increase in body fat content occurs in both γ^mt/mt^ and γ^+/+^ mice given 2% saline. Finally, these studies illustrate the utility of quantitative magnetic resonance in assessing fluid volume in dietary and genetic mouse models.

γ^mt/mt^ mice should prove valuable for further investigations into the physiologic role of ENaC’s γ subunit, an essential protein. Floxed models allow targeted deletion in specific tissues, but depend upon available Cre-expressing mouse lines (21, 36). A β subunit hypomorph with a β subunit Liddle syndrome variant has previously been reported (29, 39). The contribution of different subunits to the physiology of various cell-types likely differs, and this mouse will facilitate investigation of the physiologic role of the γ subunit.

The γ^mt/mt^ mice exhibit significantly higher aldosterone on a high K^+^ diet, suggesting impaired K^+^ secretion. In the aldosterone-sensitive distal nephron, ENaC-mediated Na^+^ reabsorption is tightly linked to K^+^ excretion. Pharmacologic or genetic impairment of ENaC activity reduce K^+^ secretion and promote hyperkalemia (4, 6). It is therefore surprising that γ^mt/mt^ mice exhibit blood K^+^ levels similar to controls. On a high K^+^ diet, the renal tubule increases K^+^ secretion through pathways that are both ENaC-dependent and ENaC-independent (13). This adaptation includes up-regulation of ROMK channels in principal cells, and BK channels in intercalated cells (7, 33). The γ^mt/mt^ mice may compensate for impaired ENaC-dependent K^+^ secretion by upregulating alternative K^+^ secretory pathways.

The γ^mt/mt^ animals exhibit no evidence of body fluid depletion at baseline, as indicated by: i) aldosterone and ANP levels that are indistinguishable from controls; ii) hemoglobin levels that reveal no signs of hemoconcentration; iii) BUN levels that are not elevated; and iv) body water and lean tissue mass that do not differ from controls. These findings stand in contrast to the induced γ subunit knock-out in the kidney of adult mice, which causes weight loss, hemoconcentration, and elevated aldosterone (5). Residual ENaC activity likely contributes, but additional compensatory mechanisms are likely. A downregulation of ENaC in non-epithelial sites (see below) might also contribute.

Dietary Na^+^ restriction produces a more rapid decline in body water and lean tissue in γ^mt/mt^ mice than in controls. More surprising is the response of γ^mt/mt^ mice given saline as drinking fluid. We predicted that impaired enteric and renal Na^+^ absorption would attenuate increases in body water associated with increased Na^+^ consumption. Instead, saline causes a significant, transient increase in total body fluid in γ^mt/mt^ animals but not controls. If confirmed, these findings may point to a role for ENaC’s γ subunit not only in promoting Na^+^ retention in the context of dietary Na^+^ depletion, but also in preventing volume-overload in the context of increased Na^+^ consumption.

Such a role may depend upon the function of ENaC in tissues beyond the kidney tubule. One group observed that global β subunit suppression increases blood pressure, as measured with intravascular telemetry catheters (9). Another group found no change in blood pressure, although they measured blood pressures in anesthetized mice using physically coupled catheters (29). Increased blood pressure and our finding of increased fluid volume sensitivity to dietary saline might reflect ENaC activity in extra-epithelial tissues where the channel is expressed, such as arterial baroreceptors, vascular endothelium and myocytes, dendritic cells, or central nervous system osmosensors (8, 10, 23, 41). Providing mice with saline to drink increases Na^+^ consumption, but also deprives them of electrolyte-free water. Further studies will be required to confirm these findings and to determine whether they are the result of fluid volume overload or relative water deprivation and increased serum tonicity.

A recent report demonstrated increased lipolysis in the context of replacement of drinking water with 2% saline (20, 30). The authors observed increased fat catabolism when dietary calorie in-take was fixed. However, the authors also noted that mice given 2% saline to drink and *ad libitum* food experience weight gain (20). As our mice were also provided food *ad libitum*, our observations are consistent with the previously reported increase in body weight, and show that the increase in weight with 2% saline specifically reflects an increase in fat mass. This illustrates an important caveat relating to the conclusion that increased salt intake stimulates lipolysis. Fat break-down may only occur when increased calorie consumption is prevented.

Finally, this study illustrates the usefulness of quantitative magnetic resonance (QMR) in evaluating body fluid content in genetic mouse models. Though percent changes in total body water were small, they likely under-represent change in extracellular water, which contributes only 1/3 of total body water (15). QMR has been used extensively for evaluation of body-fat content, but a recent study showed changes in body water due water deprivation or aldosterone administration (24). The appeal of this method stems from the ability to perform repeated measures in live, unanesthetized animals. Measurements detect changes not detectable by weight alone.

The findings in this manuscript demonstrate a novel genetic tool for exploring the importance of ENaC’s γ subunit *in vivo*. These findings, if confirmed, would suggest a surprising role for ENaC in salt-sensitivity, protecting against changes in body volume due to acute dietary increases in Na^+^ consumption.

## Acknowledgments

This work was supported by grants from the National institutes of Health, including K08 DK110332, R01 HL147818, T32 DK061296 and P30 DK079307.

## REFERENCES

1. Bagchi RA, Ferguson BS, Stratton MS, Hu T, Cavasin MA, Sun L, Lin YH, Liu D, Londono P, Song K, Pino MF, Sparks LM, Smith SR, Scherer PE, Collins S, Seto E, and McKinsey TA. HDAC11 suppresses the thermogenic program of adipose tissue via BRD2. JCI insight 3: 2018.

2. Barker PM, Nguyen MS, Gatzy JT, Grubb B, Norman H, Hummler E, Rossier B, Boucher RC, and Koller B. Role of gammaENaC subunit in lung liquid clearance and electrolyte balance in newborn mice. Insights into perinatal adaptation and pseudohypoaldosteronism. The Journal of clinical investigation 102: 1634–1640, 1998.

3. Benjamini Y, Krieger AM, and Yekutieli D. Adaptive linear step-up procedures that control the false discovery rate. Biometrika 93: 491–507, 2006.

4. Boscardin E, Perrier R, Sergi C, Maillard M, Loffing J, Loffing-Cueni D, Koesters R, Rossier BC, and Hummler E. Severe hyperkalemia is rescued by low-potassium diet in renal βENaC-deficient mice. Pflügers Archiv-European Journal of Physiology 469: 1387–1399, 2017.

5. Boscardin E, Perrier R, Sergi C, Maillard MP, Loffing J, Loffing-Cueni D, Koesters R, Rossier BC, and Hummler E. Plasma potassium determines NCC abundance in adult kidney-specific γENaC knockout. Journal of the American Society of Nephrology 29: 977–990, 2018.

6. Boyd-Shiwarski CR, Weaver CJ, Beacham RT, Shiwarski DJ, Connolly KA, Nkashama LJ, Mutchler SM, Griffiths SE, Knoell SA, and Sebastiani RS. Effects of extreme potassium stress on blood pressure and renal tubular sodium transport. American Journal of Physiology-Renal Physiology 318: F1341–F1356, 2020.

7. Carrisoza-Gaytan R, Ray EC, Flores D, Marciszyn AL, Wu P, Liu L, Subramanya AR, Wang W, Sheng S, Nkashama LJ, Chen J, Jackson EK, Mutchler SM, Heja S, Kohan DE, Satlin LM, and Kleyman TR. Intercalated cell BKα subunit is required for flow-induced K+ secretion. JCI insight 5: 2020.

8. Chung W-S, Weissman JL, Farley J, and Drummond HA. βENaC is required for whole cell mechanically gated currents in renal vascular smooth muscle cells. American Journal of Physiology-Renal Physiology 304: F1428–F1437, 2013.

9. Drummond HA, Grifoni SC, Abu-Zaid A, Gousset M, Chiposi R, Barnard JM, Murphey B, and Stec DE. Renal inflammation and elevated blood pressure in a mouse model of reduced β-ENaC. American Journal of Physiology-Renal Physiology 301: F443–F449, 2011.

10. Drummond HA, Price MP, Welsh MJ, and Abboud FM. A molecular component of the arterial baroreceptor mechanotransducer. Neuron 21: 1435–1441, 1998.

11. Drummond HA, and Stec DE. βENaC acts as a mechanosensor in renal vascular smooth muscle cells that contributes to renal myogenic blood flow regulation, protection from renal injury and hypertension. Journal of nephrology research 1: 1–9, 2015.

12. Flurkey K, and Currer JM. The Jackson Laboratory handbook on genetically standardized mice. Jackson Laboratory, 2009.

13. Frindt G, and Palmer LG. K+ secretion in the rat kidney: Na+ channel-dependent and - independent mechanisms. American journal of physiology Renal physiology 297: F389–396, 2009.

14. Grifoni SC, Chiposi R, McKey SE, Ryan MJ, and Drummond HA. Altered whole kidney blood flow autoregulation in a mouse model of reduced beta-ENaC. American journal of physiology Renal physiology 298: F285–292, 2010.

15. Hall JE, and Hall ME. Guyton and Hall textbook of medical physiology e-Book. Elsevier Health Sciences, 2020.

16. Hummler E. Epithelial sodium channel, salt intake, and hypertension. Current hypertension reports 5: 11–18, 2003.

17. Hummler E, Barker P, Gatzy J, Beermann F, Verdumo C, Schmidt A, Boucher R, and Rossier BC. Early death due to defective neonatal lung liquid clearance in alpha-ENaC-deficient mice. Nat Genet 12: 325–328, 1996.

18. Kim J-Y, van de Wall E, Laplante M, Azzara A, Trujillo ME, Hofmann SM, Schraw T, Durand JL, Li H, Li G, Jelicks LA, Mehler MF, Hui DY, Deshaies Y, Shulman GI, Schwartz GJ, and Scherer PE. Obesity-associated improvements in metabolic profile through expansion of adipose tissue. The Journal of clinical investigation 117: 2621–2637, 2007.

19. Kinsman B, Cowles J, Lay J, Simmonds SS, Browning KN, and Stocker SD. Osmoregulatory thirst in mice lacking the transient receptor potential vanilloid type 1 (TRPV1) and/or type 4 (TRPV4) receptor. American Journal of Physiology-Regulatory, Integrative and Comparative Physiology 307: R1092–R1100, 2014.

20. Kitada K, Daub S, Zhang Y, Klein JD, Nakano D, Pedchenko T, Lantier L, LaRocque LM, Marton A, and Neubert P. High salt intake reprioritizes osmolyte and energy metabolism for body fluid conservation. The Journal of clinical investigation 127: 1944–1959, 2017.

21. Malnic G, Giebisch G, Muto S, Wang W, Bailey MA, and Satlin LM. Regulation of K+ excretion. In: Seldin and Giebisch’s The KidneyElsevier, 2013, p. 1659–1715.

22. McDonald FJ, Yang B, Hrstka RF, Drummond HA, Tarr DE, McCray PB, Jr., Stokes JB, Welsh MJ, and Williamson RA. Disruption of the beta subunit of the epithelial Na+ channel in mice: hyperkalemia and neonatal death associated with a pseudohypoaldosteronism phenotype. Proceedings of the National Academy of Sciences of the United States of America 96: 1727–1731, 1999.

23. Miller RL, Wang MH, Gray PA, Salkoff LB, and Loewy AD. ENaC-expressing neurons in the sensory circumventricular organs become c-Fos activated following systemic sodium changes. American Journal of Physiology-Regulatory, Integrative and Comparative Physiology 305: R1141–R1152, 2013.

24. Morla L, Shore O, Lynch IJ, Merritt ME, and Wingo CS. A non-invasive method to study evolution of extracellular fluid volume in mice using time domain nuclear magnetic resonance. American Journal of Physiology-Renal Physiology 2020.

25. Pavlov TS, and Staruschenko A. Involvement of ENaC in the development of salt-sensitive hypertension. American Journal of Physiology-Renal Physiology 313: F135–F140, 2017.

26. Perrier R, Boscardin E, Malsure S, Sergi C, Maillard MP, Loffing J, Loffing-Cueni D, Sørensen MV, Koesters R, and Rossier BC. Severe salt–losing syndrome and hyperkalemia induced by adult nephron–specific knockout of the epithelial sodium channel α-subunit. Journal of the American Society of Nephrology 27: 2309–2318, 2016.

27. Pham CT, MacIvor DM, Hug BA, Heusel JW, and Ley TJ. Long-range disruption of gene expression by a selectable marker cassette. Proceedings of the National Academy of Sciences of the United States of America 93: 13090–13095, 1996.

28. Pitzer AL, Van Beusecum JP, Kleyman TR, and Kirabo A. ENaC in Salt-Sensitive Hypertension: Kidney and Beyond. Current Hypertension Reports 22: 69, 2020.

29. Pradervand S, Barker PM, Wang Q, Ernst SA, Beermann F, Grubb BR, Burnier M, Schmidt A, Bindels RJ, Gatzy JT, Rossier BC, and Hummler E. Salt restriction induces pseudohypoaldosteronism type 1 in mice expressing low levels of the beta-subunit of the amiloride-sensitive epithelial sodium channel. Proceedings of the National Academy of Sciences of the United States of America 96: 1732–1737, 1999.

30. Rakova N, Juttner K, Dahlmann A, Schroder A, Linz P, Kopp C, Rauh M, Goller U, Beck L, Agureev A, Vassilieva G, Lenkova L, Johannes B, Wabel P, Moissl U, Vienken J, Gerzer R, Eckardt KU, Muller DN, Kirsch K, Morukov B, Luft FC, and Titze J. Long-term space flight simulation reveals infradian rhythmicity in human Na(+) balance. Cell Metab 17: 125–131, 2013.

31. Randrianarison N, Clerici C, Ferreira C, Fontayne A, Pradervand S, Fowler-Jaeger N, Hummler E, Rossier BC, and Planes C. Low expression of the beta-ENaC subunit impairs lung fluid clearance in the mouse. Am J Physiol Lung Cell Mol Physiol 294: L409–416, 2008.

32. Ray EC. ENaC blockade in proteinuria-associated extracellular fluid volume overload - effective but risky. Physiol Rep 6: e13835, 2018.

33. Ray EC, Carrisoza-Gaytan R, Al-Bataineh M, Marciszyn A, Nkashama LJ, Chen J, Winfrey A, Flores D, Wu P, Wang W, Huang CL, Subramanya AR, Kleyman TR, and Satlin LM. L-WNK1 is required for BK channel activation in intercalated cells. AJP-Renal Physiol, under review 2020.

34. Ray EC, Chen J, Kelly TN, He J, Hamm LL, Gu D, Shimmin LC, Hixson JE, Rao DC, Sheng S, and Kleyman TR. Human epithelial Na+ channel missense variants identified in the GenSalt study alter channel activity. American journal of physiology Renal physiology 311: F908–f914, 2016.

35. Ray EC, Rondon-Berrios H, Boyd CR, and Kleyman TR. Sodium retention and volume expansion in nephrotic syndrome: implications for hypertension. Advances in chronic kidney disease 22: 179–184, 2015.

36. Rubera I, Loffing J, Palmer LG, Frindt G, Fowler-Jaeger N, Sauter D, Carroll T, McMahon A, Hummler E, and Rossier BC. Collecting duct–specific gene inactivation of αENaC in the mouse kidney does not impair sodium and potassium balance. The Journal of clinical investigation 112: 554–565, 2003.

37. Schild L. The ENaC channel as the primary determinant of two human diseases: Liddle syndrome and pseudohypoaldosteronism. Nephrologie 17: 395–400, 1996.

38. Šidák Z. Rectangular Confidence Regions for the Means of Multivariate Normal Distributions. Journal of the American Statistical Association 62: 626–633, 1967.

39. Snyder PM. The epithelial Na+ channel: cell surface insertion and retrieval in Na+ homeostasis and hypertension. Endocr Rev 23: 258–275, 2002.

40. Taicher GZ, Tinsley FC, Reiderman A, and Heiman ML. Quantitative magnetic resonance (QMR) method for bone and whole-body-composition analysis. Analytical and bioanalytical chemistry 377: 990–1002, 2003.

41. Tarjus A, Maase M, Jeggle P, Martinez-Martinez E, Fassot C, Loufrani L, Henrion D, Hansen PB, Kusche-Vihrog K, and Jaisser F. The endothelial αENaC contributes to vascular endothelial function in vivo. PloS one 12: e0185319, 2017.

42. Theander-Carrillo C, Wiedmer P, Cettour-Rose P, Nogueiras R, Perez-Tilve D, Pfluger P, Castaneda TR, Muzzin P, Schürmann A, Szanto I, Tschöp MH, and Rohner-Jeanrenaud F. Ghrelin action in the brain controls adipocyte metabolism. The Journal of clinical investigation 116: 1983–1993, 2006.

43. Tinsley FC, Taicher GZ, and Heiman ML. Evaluation of a Quantitative Magnetic Resonance Method for Mouse Whole Body Composition Analysis. Obesity Research 12: 150–160, 2004.

44. Unruh ML, Pankratz MS, Demko JE, Ray EC, Hughey RP, and Kleyman TR. Trial of Amiloride in Type 2 Diabetes with Proteinuria. Kidney Int Rep In press, 2017.

